# MAAD: Multidimensional Antiviral Antibody Database

**DOI:** 10.1101/2025.11.18.688681

**Authors:** Yixin Li, Jinyue Wang, Chuziyue Zhang, Yuxia Zhang, Jing Deng, Han Zhang, Mingkai Li, Fan Wang, Xiangxi Wang

## Abstract

Antibodies have emerged as central components of therapeutic strategies against viral infectious diseases, functioning as key effectors in both prevention and treatment. While traditional antibody discovery has relied heavily on high-throughput screening, the field is now shifting toward rational antibody design, which requires integrative insights into sequence-structure-function relationships. However, the absence of a standardized and well-annotated antibody database integrating these multidimensional features hampers systematic exploration, cross-pathogen comparison, and rational antibody design. Here, we introduce a “Multidimensional Antiviral Antibody Database” (MAAD; http://www.xxx), a curated platform dedicated to antibody nanobody and single-chain variable fragment targeting three high-impact RNA virus families, Coronaviridae (SARS-CoV-1, SARS-CoV-2, MERS-CoV), Orthomyxoviridae (influenza virus), and Pneumoviridae (respiratory syncytial virus, human metapneumovirus), due to the large, high-quality datasets accumulated in recent years. MAAD further incorporates a suite of interactive analysis modules, including CDR annotation, similarity-based CDR3 sequence analysis, V/J gene usage profiling, sequence-based clustering and structure-based antigen-antibody interfaces residues with per-site entropy and mutation rate profiling. These features enable in-depth exploration of antibody sequence characteristics, thereby facilitating functional and structural insights for rational antibody design. Together, by bridging antibody sequence, structure and function, MAAD offers an open and standardized platform that advances comparative antiviral research and supports therapeutic antibody discovery.

## Introduction

Since the development of hybridoma technology, which enabled the generation of monoclonal antibodies (mAbs), mAbs have emerged as one of the most important classes of biotherapeutics, not only for the treatment of oncologic and autoimmune diseases, but also for combating viral infectious diseases (Köhler and Milstein 1975; Yasunaga 2020; Pantaleo et al. 2022; Paul et al. 2024). In particular, mAbs are promising prophylactic and therapeutic agents for viral infections due to their high specificity and immune-enhancing properties. Several antiviral mAbs have been approved by the U.S. Food and Drug Administration (FDA). For example, palivizumab was the first FDA-approved mAb for the prevention of respiratory syncytial virus (RSV) infection (Young 2002); ibalizumab was authorized for the treatment of HIV-1 infection (Markham 2018); and ansuvimab received approval in 2020 for the treatment of Ebola virus (EBOV) infection (Lee 2021). During the COVID-19 pandemic, several mAbs such as sotrovimab, casirivimab and bamlanivimab, received Emergency Use Authorization (EUA) from the FDA (Deeks 2021; Heo 2022). While vaccines have played a central role in controlling the COVID-19 pandemic, mAbs have served as a vital countermeasure for high-risk populations, such as immunocompromised individuals, thereby underscoring their critical role in mitigating emerging viral threats (Schmidt et al. 2024).

Building on these clinical advances, attention has increasingly shifted toward efficient strategies to accelerate mAbs discovery and optimization. Traditional antibody discovery has relied heavily on high-throughput screening (Mahdavi et al. 2022). However, the field is now rapidly shifting toward rational antibody design, which requires a comprehensive and deep understanding of sequence, structure and function relationships. Importantly, this understanding not only enables the engineering of antibodies with desired properties, such as enhanced specificity or cross-reactivity, but also informs antibody-based vaccinology (Lanzavecchia et al. 2016). Specifically, antibody-based vaccinology aims to overcome the limitations of traditional vaccine approaches by designing novel immunogens based on structural characterization of antigen-antibody complexes (Pantaleo et al. 2022). By identifying protective epitopes and masking immunodominant but non-neutralizing regions, this approach focuses the immune response on functionally critical targets, thereby enhancing vaccine efficacy. This strategy has been successfully applied in the design of immunogens targeting conserved neutralizing epitopes on influenza hemagglutinin (HA) (Weidenbacher and Kim 2019). It has also been utilized in vaccines that display SARS-CoV-2 receptor-binding domains (RBDs) in a highly immunogenic array and exhibit a lower antibody binding:neutralizing ratio (Walls et al. 2020). The above-mentioned successes highlight the dual role of mAbs as both therapeutic agents and blueprints for vaccine design. Meanwhile, to address the growing demand for mAbs discovery, artificial intelligence (AI) has emerged as a powerful tool to accelerate their identification and optimization (Lou et al. 2023). However, the performance of AI-driven approaches critically depends on large-scale, standardized training data that systematically connect sequences and structures to their functional properties. To address the growing demand for rational antibody design, antibody-based vaccinology and AI-driven antibody discovery, there is an urgent need for a standardized, well-annotated antibody database that integrates sequence, structure and function data into a coherent platform.

One of the major challenges in antibody engineering is to elucidate the relationships among sequence, structure and function that govern antibody specificity and breadth. Although existing antibody databases have provided valuable resources to the field (Dunbar et al. 2014; Raybould et al. 2021; Olsen et al. 2022), there remains a need for a platform that comprehensively integrates sequence, structural and functional annotations across diverse viral pathogens (Table. 1). To address this gap, we here introduce a multidimensional database of antiviral antibody (MAAD) which integrates 27,707 antibody, nanobody and single-chain variable fragment (scFv) entries (Fig. 1 and S1A). MAAD focuses on antibodies targeting three high-impact RNA virus families including Coronaviridae (Severe Acute Respiratory Syndrome Coronavirus 1 [SARS-CoV-1], Severe Acute Respiratory Syndrome Coronavirus 2 [SARS-CoV-2], Middle East Respiratory Syndrome coronavirus [MERS-CoV]), Orthomyxoviridae (influenza virus), and Pneumoviridae (respiratory syncytial virus [RSV], human metapneumovirus [hMPV]), due to the large, high-quality datasets accumulated in recent years (Fig. 2A). These antibodies benefit from standardized binding and neutralization assays, making them ideal for building a comprehensive, standardized, functionally annotated database.

**Figure 1.**
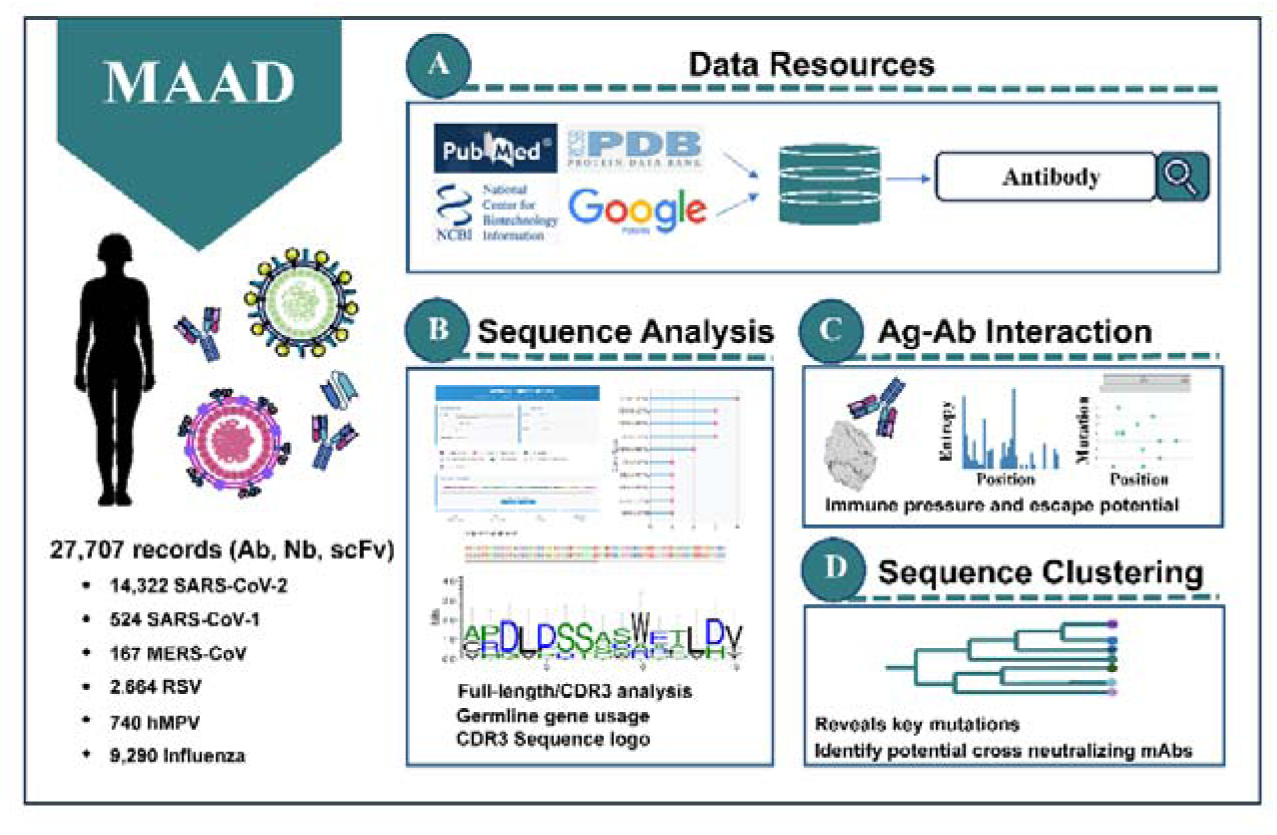
Overview of MAAD and its functional modules. (A) Antibody, nanobody and scFv entries were systematically collected from multiple sources including peer-reviewed publications, patents, and clinical sources, and integrated into the unified MAAD database with standardized annotations. (B) The platform supports interactive sequence analysis, including CDR annotation, V/J germline gene assignment, and similarity-based searches using full-length or CDR sequences. (C) For antibodies with resolved structures, MAAD provides detailed interface residue annotations, coupled with per-site Shannon entropy and mutation frequency based on viral sequence diversity. (D) A sequence-based clustering and phylogenetic tree construction module implemented to group antibody/nanobody.

**Figure 2.**
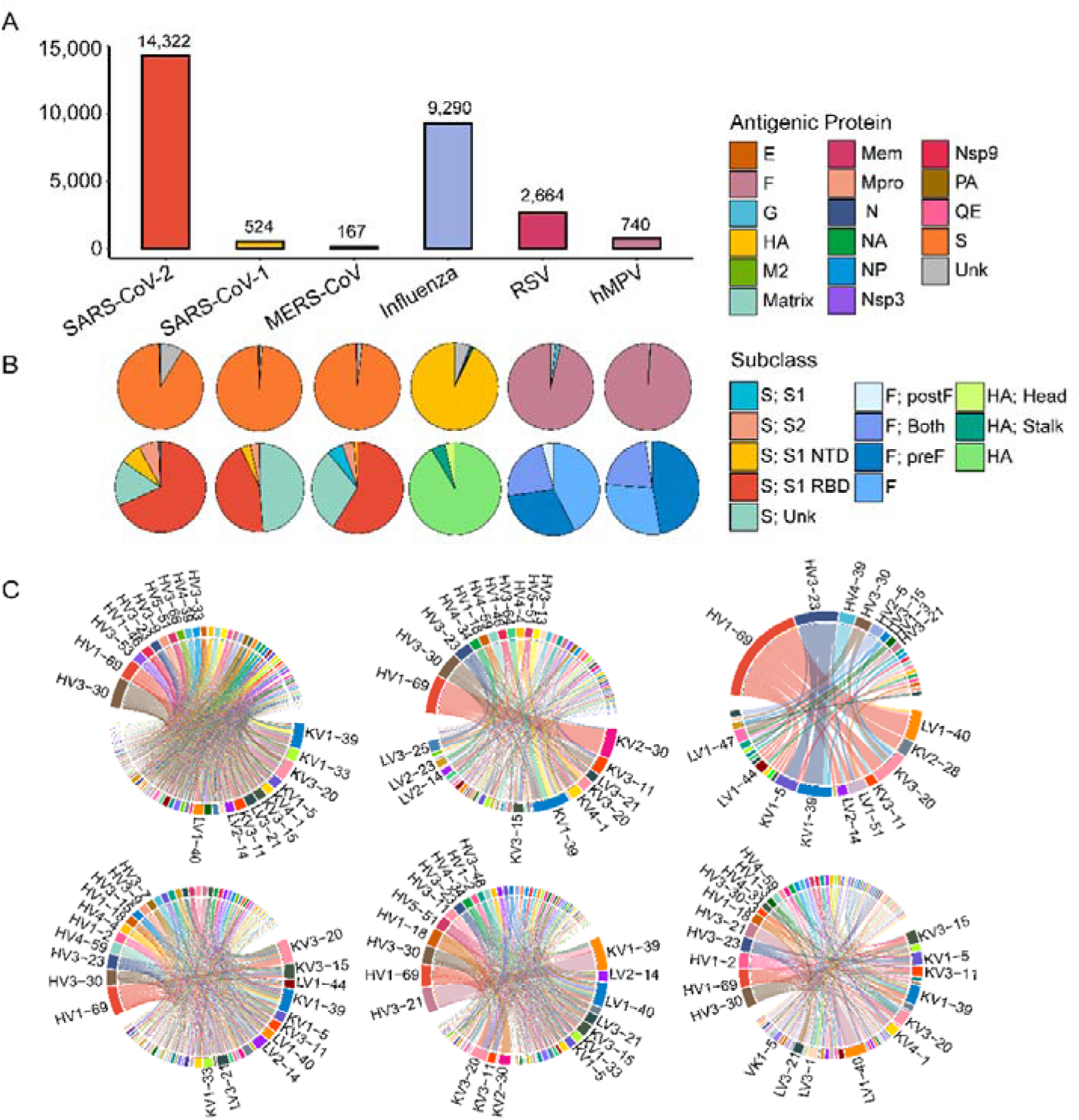
MAAD statistics and antibody gene usage profiles. (A) Number of curated entries primarily derived from three viral families including *Coronaviridae* (SARS-CoV-1, SARS-CoV-2, MERS-CoV), *Orthomyxoviridae* (influenza A/B), and *Pneumoviridae* (RSV, hMPV). (B) Pie chart of the first row illustrating the distribution of antigenic protein targets. S = Spike protein; N = Nucelocapsid Protein; Mem = Membrane Protein; E = Envelope Protein; Nsp = Non-structural Protein; Mpro = Main Protease; HA = Hemagglutinin; NA = Neuraminidase; NP = Nucleoprotein; M2 = Matrix Protein 2; F = Fusion Protein; G = Attachment protein; QE = Quaternary epitope; Unk = Unknown. Pie chart of second row illustrating the distribution of dominant antigenic protein subcategories across different viruses. For coronaviruses (SARS-CoV-1, SARS-CoV-2, and MERS-CoV), spike-specific antibodies are further classified as: S1, S2, S1-RBD (Receptor Binding Domain), S1-NTD (N-terminal RNA-binding domain), and unknown regions. “S1” refers to antibodies mapped to the S1 subunit but lacking finer resolution to assign them to RBD or NTD; “Unknown” denotes antibodies known to bind S protein but with no specific subunit (S1/S2) information available. For RSV and hMPV, the F protein is categorized into prefusion (pre-F), postfusion (post-F), or cross-binding to both. For influenza viruses, hemagglutinin (HA) specific antibodies are subdivided into those targeting the HA head or stalk regions. (C) Circos plot showing the top 10 IGHV-IGLV pairings observed across antigen-specific human derived antibodies (only gene pairs with frequency >1 shown).

To further facilitate user-driven analysis, MAAD also integrates a suite of interactive analysis and visualization modules for complementarity determining regions (CDRs) annotation and similarity-based entry search, V/J gene usage profiling, identification of antigen-antibody interface residues, per-site Shannon entropy and mutation analysis, as well as phylogenetic clustering of antibody sequences. By linking antibody sequence, structure, and function through integrated analysis and visualization modules, MAAD provides not only a valuable platform for rational antibody and vaccine design, but also a robust data foundation for training AI-driven models in paratope prediction, cross-reactivity assessment, and antiviral antibody discovery.

## Results

### Comprehensive functional annotation of entries in MAAD

Currently, MAAD integrates 27,707 standardized antibody, nanobody and scFv entries compiled from 805 peer-reviewed publications and 140 patents. These entries cover six high-impact respiratory pathogens across three viral families, including *Coronaviridae* (SARS-CoV-1, SARS-CoV-2, MERS-CoV), *Orthomyxoviridae* (influenza A and B), and *Pneumoviridae* (RSV, hMPV) (Fig. 2A). Among the 27,707 entries, approximately 17,700 entries are experimentally annotated with detailed functional data such as binding and/or neutralization. The remaining entries, although lacking direct functional annotations, are primarily derived from antigen-specific B cell repertoires. These sequence-only entries retain complete variable region sequences and serve as a valuable resource for clonal lineage inference and the training of machine learning models for antibody function prediction.

Each entry is annotated with key metadata, including the published name of the antibody, nanobody and scFv, antigen target, biological or synthetic origin (e.g., infected human, immunized mouse, engineered) and the experimentally validated antigen-binding and/or neutralization specificities (Table S1). Full-length variable region sequences of each entry are numbered using ANARCI (Dunbar and Deane 2016), which employs Hidden Markov Models to align input sequences to pre-numbered germline references. CDRs are annotated based on three standardized numbering schemes: the international IMGT (Lefranc et al. 2003), Kabat (Kabat and Wu 1971) and Chothia (Chothia and Lesk 1987). When available, nucleotide sequences are included, along with corresponding GenBank (Benson et al. 2011) accession numbers. Structural information is linked directly to corresponding Protein Data Bank (PDB) (Berman et al. 2000) entries. In addition, all records are cross-referenced to their original literature sources with PubMed ID (PMID) or patent number and publication date.

Specifically, the majority of entries are directed against SARS-CoV-2 (WT=12,077; Alpha=735; Beta=1,163; Gamma=735; Delta=1,191; Epsilon=2,471; Omicron=3,125), reflecting the intensive research focus during the COVID-19 pandemic. The database additionally comprises entries targeting MERS-CoV (n=70), SARS-CoV-1 (n=2,209), influenza viruses (influenza A=1,475; influenza B=248), RSV (RSV-A=1,478; RSV-B=907), and hMPV (hMPV-A=290; hMPV-B=270) (Fig. S1B). Notably, entries are not mutually exclusive across antigens, as individual antibodies may have been tested against multiple targets. Based on all antibody and nanobody entries collected in the MAAD database, we analyzed the distribution of viral protein targets (Fig. 2B). For all three coronaviruses (SARS-CoV-1, SARS-CoV-2, and MERS-CoV), the spike protein is the predominant target, with the RBD being the most investigated domain (Fig. 2B). In the case of influenza virus, HA is the predominant target of antibody binding, accounting for over 93% of entries, while antibodies against other viral components such as nucleoprotein (NP) and neuraminidase (NA) are extremely rare (<1%) (Fig. 2B). For Pneumoviridae members RSV and hMPV, most antibodies are directed against the fusion (F) protein. For RSV, over 96% of antibodies bind to F protein, with approximately 2% targeting the attachment glycoprotein (G) (Fig. 2B). In hMPV, the vast majority of antibodies (99%) target the F protein, whereas only a small fraction recognizes the matrix protein (Fig. 2B). Notably, we observed that for both RSV and hMPV, F-specific antibodies mainly recognize the pre-fusion conformation or exhibit cross-reactivity, whereas post-fusion specific antibodies are rare (Fig. 2B). MAAD also incorporates 67 clinically evaluated therapeutic antibodies from regulatory documents and published clinical studies. These entries cover antibodies targeting viral antigens represented in MAAD, including SARS-CoV-2, MERS-CoV, RSV and influenza, thereby providing important benchmarks for therapeutic development. By including antibodies that have advanced into clinical use, MAAD offers real-world evidence of efficacy and safety, which enables comparative analyses with preclinical candidates.

### Interactive exploration and visualization for antibody sequence analysis

To facilitate intuitive exploration and functional analysis of antibody and nanobody entries, MAAD implements a set of interactive modules for data query, visualization, and sequence analysis. Users can perform name-based searches to quickly locate specific antibody or nanobody entries of interest, or use virus-based searches to identify entries derived from B cells exposed to a particular virus through infection or immunization. Additional filters include developmental origin (e.g., human, murine, camelid, synthetic), V/J germline gene usage, PDB and project-specific identifiers (Fig. S4A). Each matched entry links to a detailed information page including comprehensive metadata such as targeted epitopes, full-length variable region sequences, CDR annotations across IMGT, Kabat, and Chothia schemes, structural information (PDB), and available binding or neutralization profiles. Meanwhile, references are hyperlinked, allowing users to trace each record back to its original publication or patent source for straightforward verification (Fig. S4A).

In addition to metadata-based queries, MAAD also supports sequence-based similarity searches, enabling users to input custom antibody variable region sequences and retrieve matched entries with annotated V/J germline gene information (Fig. 3A and S4B). The input sequence is first processed through ANARCI and returned with annotated CDRs and corresponding V/J germline genes. Two analysis modes are available: (1) full-length sequence similarity analysis mode, where the query sequence is aligned against all MAAD entries using BLAST (Altschul et al. 1990) with a user-defined similarity threshold. Matched entries are summarized with V/J gene usage dot plots to assess germline convergence or divergence patterns (Fig. 3B). (2) CDR-focused analysis mode, where users may focus on specific CDR regions (e.g., CDR3) as input. Matched sequences of identical length or those containing the specified motif, are visualized with sequence logo plots generated by WebLogo (Crooks et al. 2004) (Fig. S4B), thereby highlighting conserved and variable positions. Associated V/J gene dot plots are also provided to reveal lineage biases within matched antibody repertoires.

**Figure 3.**
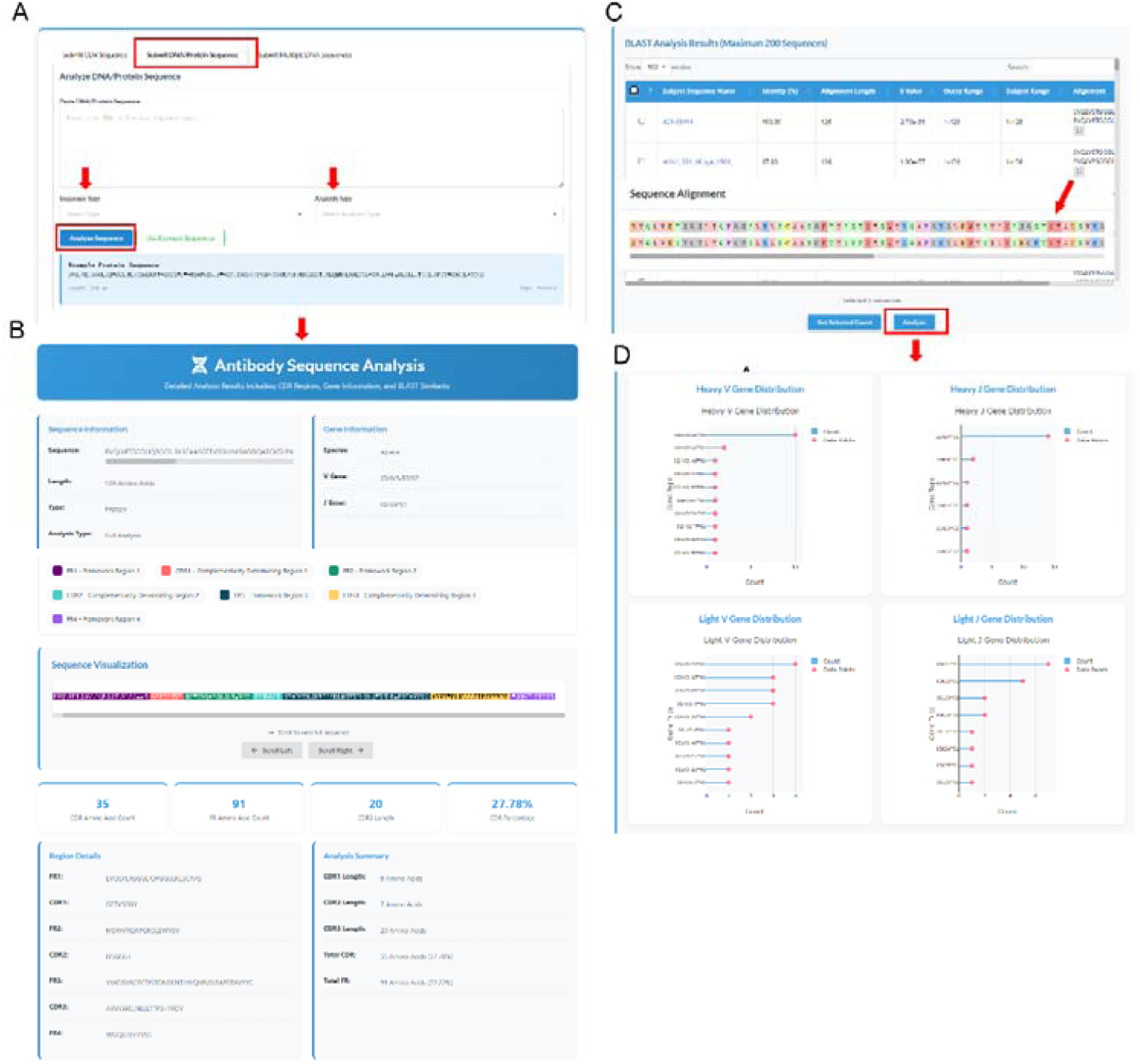
Detail page of sequence analysis model. (A) Interactive interface for sequence-based similarity search. (B) Full-length antibody sequence results with V/J germline assignment and framework/CDR annotation based on the IMGT scheme. (C) Results of sequence-based similarity searches. (D) Statistical summary of selected entries, with dot plots illustrating germline gene distributions.

To supporting interactive analysis of user-provided sequences, MAAD offers an overview of germline gene usage patterns across the entire database. By profiling V and J gene pairings in both heavy and light chains, we revealed distinct patterns across viral targets (Fig. 2C and S3). Among human derived antibodies, certain IGHV genes such as IGHV3-30 and IGHV1-69 were broadly utilized across multiple viruses, however, their light chain partners exhibited considerable diversity (Fig. 2C). For example, IGLV1-40, IGKV3-11, and IGKV3-20 frequently paired with IGHV1-69 in SARS-CoV-2 antibodies, whereas IGKV2-30 was the predominant light chain partner for IGHV1-69 in SARS-CoV-1. Notably, the IGHV1-69/IGKV3-20 pairing was highly enriched in influenza targeting antibodies and IGHV1-69/IGKV3-15 pairing dominated in hMPV-targeting responses. In contrast to the broad usage of IGHV1-69, RSV antibodies exhibited a distinct preference for IGHV3-21, highlighting virus-specific germline biases across pathogens (Fig. 2C). An overview of CDR3 length distributions is also provided. Heavy-chain CDR3s generally displayed broader variability than light chains, with light-chain CDR3 lengths concentrated around 9–11 amino acids, whereas heavy-chain CDR3s were distributed more broadly, ranging from 10 to 23 amino acids (Fig S2). Together, these analyses highlight both conserved and virus-specific features of antibody repertoires.

### Integration and annotation of antigen-antibody interaction profiles

Given the continuous emergence of viral variants, structural information is essential for dissecting antibody-antigen interactions. MAAD currently integrates 1,394 resolved antigen-antibody complex structures and each entry is linked to its corresponding PDB identifier. Detailed interaction profiles can be explored by clicking the PDB identifier, which directs users to a page specifying the antigen chain(s) and the antibody heavy and light chains (Fig. 4A). To characterize binding interfaces, interface residues were defined as antigen or antibody residues with at least one interatomic distance of less than 4.5 Å.

**Figure 4.**
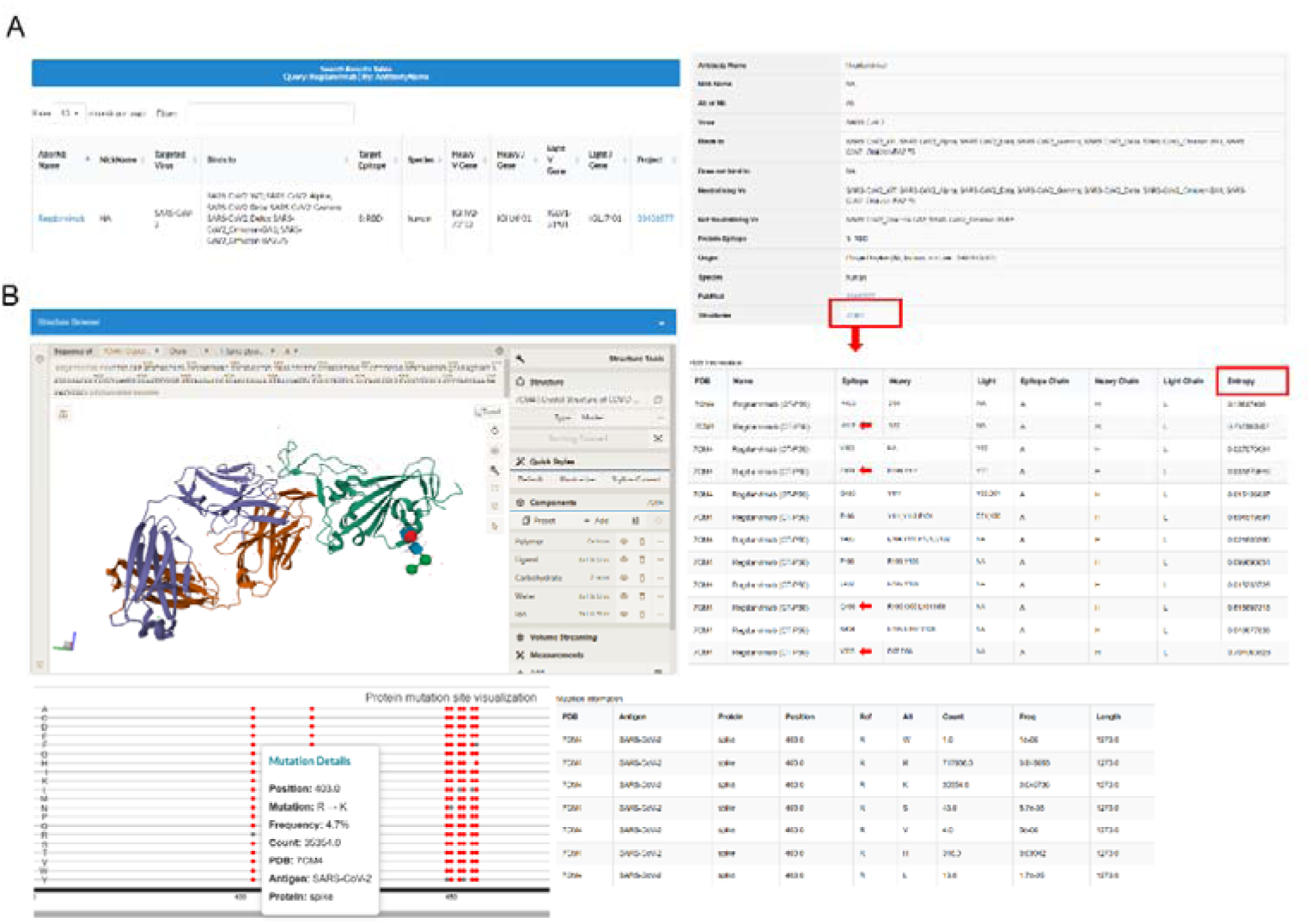
Detail page of antigen-antibody complexes interaction profiles. (A) Example workflow illustrating how resolved antigen-antibody complex structures are identified. (B) Detail page of resolved antigen-antibody structures and Shannon entropy of interface residues for each pathogen (top). Dot plot representation of mutation frequencies at interface residues across pathogens (bottom).

To assess antigenic variability at antigen-antibody interfaces, MAAD annotates interface residues with Shannon entropy scores, reflecting site-specific amino acid diversity (Fig. 5A-B and S5A-B). For coronaviruses, entropy was calculated for spike protein, which comprises the majority of resolved antigen-antibody complexes (Fig. 2B and 5A). In SARS-CoV-2, elevated entropy was observed within spike residues 340–510, corresponding to the receptor-binding domain (RBD) and receptor-binding motif (RBM), well-known hotspots of immune pressure (Jian et al. 2025). By contrast, MERS-CoV spike displayed relatively uniform entropy, while the limited number of sequences restricted the analysis of SARS-CoV-1. For RSV and hMPV, entropy and mutation rate profiles were assessed independently for the F and G proteins from both subtype A and B strains (Fig. 5B and S5B). The F protein of both RSV-A and RSV-B displayed generally low variability, however RSV-B showed pronounced peaks of entropy and mutation frequency between residues 170–210 compared with RSV-A (Fig. 5B). In contrast, the RSV G protein was markedly more variable (Fig. S5B), with both RSV-A and RSV-B exhibiting extensive entropy peaks and frequent mutations across the mucin-like regions, while the central conserved domain (CCD) remained relatively stable. For hMPV, the F protein of both subtypes also exhibited overall low variability, with sporadic peaks of entropy and mutations primarily within the F1 subunit (Fig. 5B). The hMPV G protein, similar to RSV G, is highly variable due to its mucin-like domains (Fig. S5B). These observations highlight that while both RSV and hMPV F proteins are generally conserved with subtype-specific differences in variability distribution, their G proteins exhibit extensive sequence diversity. For influenza virus, the major surface glycoprotein hemagglutinin (HA) is critical for facilitating virus entry and infection of host cells, and it exhibits relatively low sequence conservation across strains owing to its antigenic diversity and rapid evolution (Wilson et al. 1981; Wu and Wilson 2020). To investigate sequence variability, we collected HA sequences from human-derived strains of major public health concern (H1N1, H3N2, H5N1, H7N9, and two influenza B lineages) (Su et al. 2017; Fasanmi et al. 2017; Bi et al. 2024) and performed Shannon entropy analyses separately. Consistent with previous research, HA1 exhibited relatively higher entropy scores than HA2 across all examined influenza subtypes highlighting HA1 as the major target of antigenic drift (Fig. S5A). Peaks of sequence variability were predominantly concentrated within HA1, whereas HA2 showed overall lower entropy, reflecting its greater sequence conservation (Fig. S5A).

**Figure 5.**
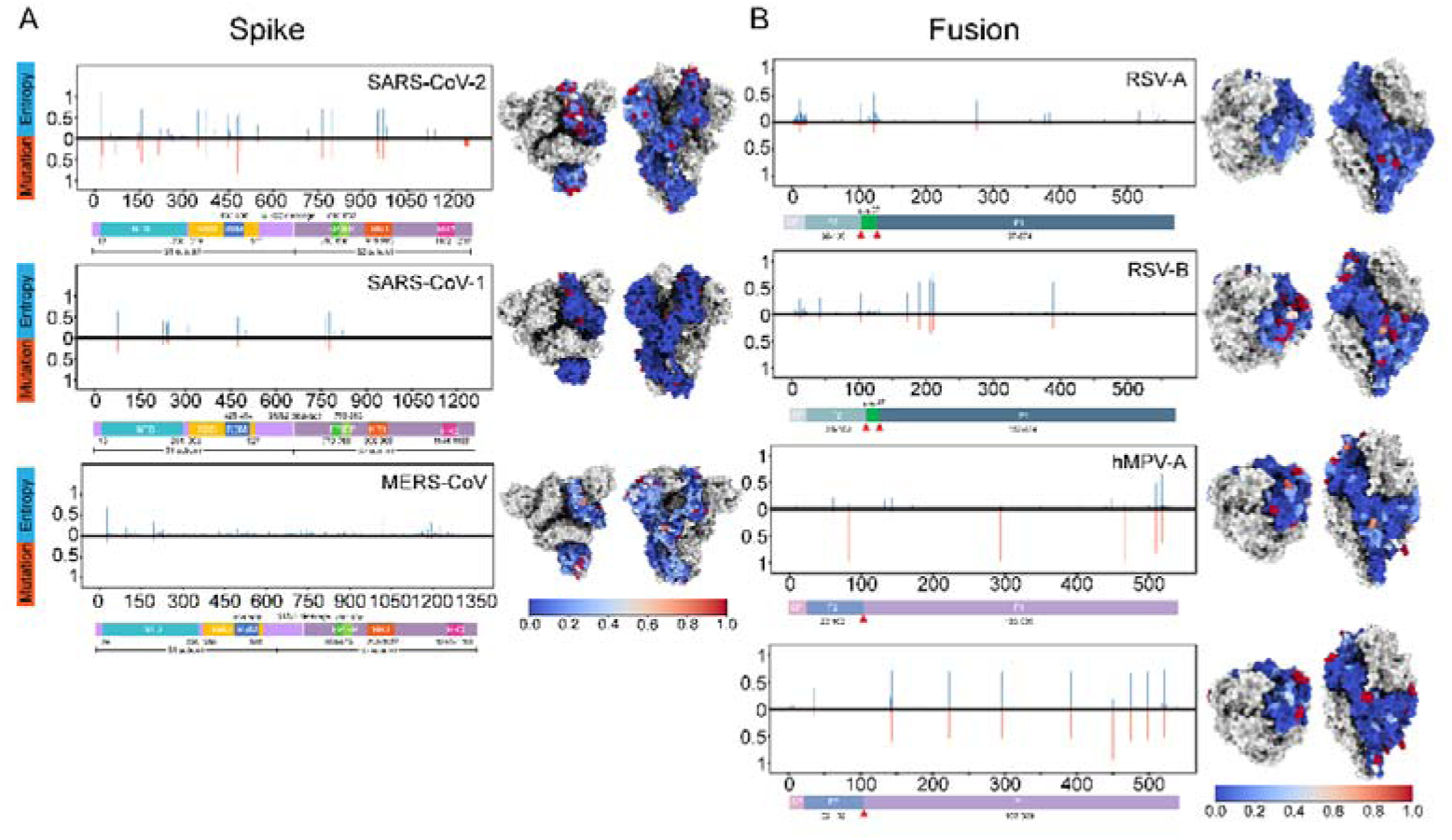
Viral entropy and mutation analysis. (A) and (B) show site-specific sequence variability and mutational landscape of viral antigens. Bar plots display per-residue Shannon entropy (blue) and mutation frequency (orange), calculated based on aligned viral sequences relative to a reference strain (A: spike protein; B: fusion protein). Corresponding 3D antigen structures colored by normalized Shannon entropy values to visualize surface variability. Two structural orientations are shown: top view (left) and side view (right).

The integrative analysis of binding interfaces and sequence-level variability facilitates the elucidation of molecular interactions and provides guidance for antibody design and optimization. For instance, although Regdanvimab exhibited neutralizing activity against multiple SARS-CoV-2 variants, including Gamma, Delta, Epsilon, and Kappa, it has been demonstrated to have significant escape from Omicron variants (Planas et al. 2022). Structural analysis of the Regdanvimab-spike complex in MAAD highlights several key contact residues that display both high entropy and high mutation frequency (Fig. 4B). Notably, mutations such as K417N, E484A, Q493R and Y505H have been proven to directly impact the antigen-antibody interface and correlate with the loss of neutralization potency (Cao et al. 2022).

### Sequence-based clustering and tree construction of antibodies

With the rapid growth of antibody repertoire data generated by next-generation sequencing (NGS), a substantial portion of NGS-derived sequences remain experimentally unvalidated, lacking direct evidence of antigen binding or neutralization. In this context, MAAD integrates a sequence-based clustering and a phylogenetic reconstruction module to cluster functionally uncharacterized antibodies alongside annotated ones, thereby facilitating comparative analysis and hypothesis generation regarding potential functional similarity (Fig. 6A-B).

**Figure 6.**
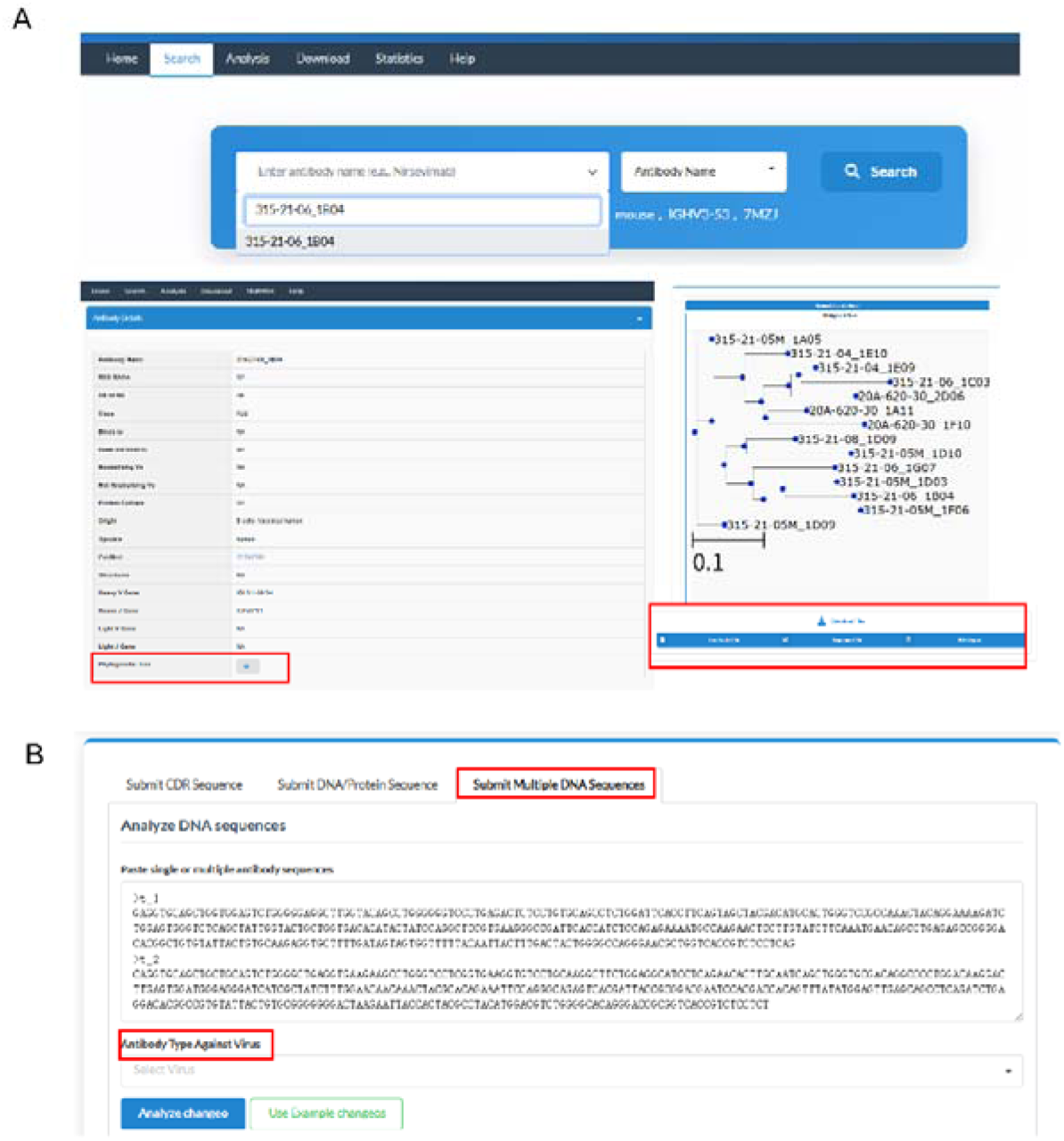
Detail page of sequence-based phylogenetic clustering and tree construction. (A) Example workflow illustrating how to view and download precomputed phylogenetic trees that integrate both functionally characterized and uncharacterized entries across the database. (B) Interactive interface for user-submitted sequence phylogenetic clustering.

This module enables users to perform clonal grouping based on V/J germline gene usage and CDR3 sequence similarity. For heavy chain sequences, clonotype assignment was inferred with the Change-O toolkit (Gupta et al. 2015), based on germline annotations to cluster similar sequences. Within each assigned cluster, sequences were aligned using MAFFT (Katoh et al. 2002), and maximum-likelihood phylogenetic trees were generated by IQ-TREE (Nguyen et al. 2015) with appropriate evolutionary models to illustrate the phylogenetic relationships among clonally related antibodies (Fig. S6A). These phylogenetic trees provide deep insights into patterns of amino acid mutation and potential functional convergence within antibody families. When integrated with functional annotations, such as antigen binding specificity or neutralization breadth, the trees enable users to identify key mutations associated with antigen recognition or track the evolution of broad neutralization capacity. For entries lacking direct functional evidence, phylogenetic proximity to well-characterized antibodies can offer indirect inferences regarding their target specificity, cross-reactivity, or therapeutic potential. For example, within a clonally related group of RSV-targeting antibodies, all members displayed binding to RSV A2 and B18537 strains. However, only a subset (cluster 2) exhibited strong neutralization activity against RSV B18537, whereas others (cluster 1) showed negligible neutralization (Fig. S6B). This highlights how phylogenetic clustering, when combined with functional data, can differentiate between binding-only and neutralizing antibodies within the same lineage, thereby uncovering mutations associated with the acquisition of neutralization breadth (Fig. S6B).

MAAD supports two modes of phylogenetic analysis: (1) user-driven phylogenetic tree reconstruction, in which user-uploaded sequences are integrated with MAAD sequences to infer a combined phylogenetic tree; and (2) exploration of precomputed phylogenetic trees that integrate both functionally characterized and uncharacterized entries across the database (Fig. 6A-B and S6A). These two modes enable investigation of the evolutionary context of antibodies and the inference of potential functions through tree-based similarity. Tree files can be retrieved by antibody name, with visualizations available in PNG format. In addition, a corresponding annotation table is also provided, presenting mutation profiles and functional properties of related antibodies in a structured format. This facilitates downstream analyses of antibody evolution under immune selection pressure.

## Discussion

In this study, we developed MAAD, a multidimensional antiviral antibody database targeting pathogens from three major RNA viral families. The current version of MAAD includes 27,707 curated entries, each annotated with standardized metadata including amino acid and nucleotide sequences, V/J germline usage, CDRs, targeted antigens, functional annotations, and structural information (Table. S1). In addition, MAAD incorporates clinically evaluated therapeutic antibodies, providing a unique benchmark for comparison and enabling translational insights derived from well-characterized and successful antiviral agents.

In parallel, MAAD also provides a suite of interactive modules to support in-depth exploration of antibody sequence features, antigen-antibody complexes interfaces, and functional inference through tree-based similarity. The platform enables CDR annotation, V/J gene usage profiling, and both full-length and CDR-based similarity searches, complemented by visual tools such as germline dot plots and sequence logo diagrams. This integrative framework reveals both conserved and virus-specific patterns of germline usage, providing intuitive insights and a systematic reference for evaluating gene biases across antiviral responses. Meanwhile, it transforms MAAD from a static repository into a dynamic, user-friendly platform for antibody repertoire profiling and the elucidation of sequence-function relationships. To complement these sequence-level analyses, MAAD also incorporates structural insights, which are essential for understanding viral immune escape. The emergence of SARS-CoV-2 variants during the COVID-19 pandemic demonstrated the virus’s ability to evade vaccine or infection induced antibodies, leading to breakthrough infections and reinfections (Carabelli et al. 2023; Zhang et al. 2024). These observations underscore the importance of elucidating molecular interactions between antibodies and viral antigens, which serve as the foundation for rational antibody therapeutic design. For antibodies with resolved antigen-antibody complexes structures, MAAD systematically maps interface residues and annotates them with site-specific Shannon entropy and mutation frequency metrics derived from aligned viral sequences, allowing assessment of immune pressure and escape potential. MAAD also supports phylogenetic clustering and tree construction for user-submitted sequences, as well as exploration of precomputed trees that integrate both functionally characterized and uncharacterized antibodies. This module is particularly relevant in the context of next-generation sequencing (NGS) of antigen-responding B cell repertoires, which generates large-scale antibody sequences but often lacks direct experimental validation (Goldstein et al. 2019). Computational clustering and phylogenetic reconstruction therefore provide a valuable means to infer antibody specificity and functional potential from uncharacterized sequences. These trees, paired with mutational and binding annotations, support functional inference, enable detailed evolutionary analyses, and facilitate the identification of promising therapeutic leads through sequence-based clustering and functional annotation. Taken together, these modules establish a comprehensive framework for understanding of antigen recognition and provide actionable guidance for rational antibody design and optimization.

Moreover, MAAD will continue to expand in order to support a broader spectrum of pathogens driven by the incorporation of new antibody data and the need to address emerging research priorities. Such expansion will bridge critical knowledge gaps and enhance the platform’s value for early-stage therapeutic discovery and pandemic preparedness. To support this growth, MAAD employs a modular and extensible architecture that enables the integration of additional data types, such as deep mutational scanning (DMS) (Fowler and Fields 2014) results and quantitative experimental measures including binding affinity (e.g., KD) and neutralizing activity (e.g., IC50). Importantly, the standardized structure of MAAD entries makes it well-suited for AI-driven applications. Its curated integration of antibody sequence, structure, and function provides a robust foundation for machine learning. Features such as aligned full-length antibody sequences, variable regions, V/J gene assignments, and mapped structural binding residues serve as high-quality input data for deep learning models. These resources create a fertile training ground for machine learning models in tasks, such as paratope prediction, neutralization classification, and cross-reactivity forecasting, all supported by experimentally validated data. Looking ahead, we plan to incorporate standardized NGS datasets from antigen-enriched B cell repertoires, particularly those with paired heavy-light chains and confirmed antigen specificity. Collectively, these enhancements will greatly extend the scope and the power of AI-driven discovery within MAAD, reinforcing its role as both a research resource and a platform for translational innovation.

In summary, MAAD serves not only as a comprehensive repository of antiviral antibody database, but also as a versatile and extensible platform for sequence-function-structure integration, thereby supporting rational therapeutic antibody design, and AI-assisted antibody discovery. Through the integration of large-scale curated data and versatile analytic platforms, MAAD provides a foundation for rational antibody design, therapeutic optimization, and AI-driven discovery, ultimately advancing preparedness against current and future viral threats.

**Table 1.** Comparison of MAAD with existing antibody databases.

| Feature | CoV-AbDab<br>(Raybould et al. 2021) | SAbDab<br>(Dunbar et al. 2014) | OAS<br>(Olsen et al. 2022) | MAAD |
| --- | --- | --- | --- | --- |
| Scope | Ab/Nb against for CoVs | Ab/Nb with resolved structures | Broad | Ab/Nb against several antigens |
| Sequence type | Paired aa & nt sequence | Paired aa & nt sequence | Unpaired and paired aa & nt sequence | Paired aa & nt sequence |
| Functional annotation | Binding, neutralization | Binding with resolved structures | No function, only repertoire | Binding, neutralization |
| Structural annotation | None | None | None | Antigen-antibody interface residues annotated |
| Virological data | None | None | None | Per-residue Shannon entropy & mutation rate at interface positions |
| Interactive modules | Basic and sequence-based search | Basic search and structural viewer | Basic search | Basic and sequence-based search, CDR annotation, germline usage profiling, antigen-antibody interfaces structural viewer, phylogenetic clustering |

\*aa: amino acid, nt: nucleotide, Ab/Nb: antibody/nanobody

## Acknowledgments

We gratefully acknowledge all the authors of the studies listed in the database metadata for providing the underlying data that formed the basis of this work. Their contributions in generating and sharing these experimental data have been invaluable to our analyses. Additionally, we extend our thanks to the group members and the many users who reported bugs and offered constructive suggestions, that greatly improved the quality and usability of the database.

## Author contributions

XX.W, JY.W, and F.W designed and supervised the study. JY.W and YX.L conceived and designed the study. YX.L conducted the investigation, curated the data, developed the analysis pipeline and scripts, carried out the methodology and formal analysis, and drafted the original manuscript. JY.W contributed to the investigation, carried out the methodology and constructed the database. CZY.Z contributed to the investigation and data curation. J.D provided the scripts for tree visualization. MK.L curated the data. YX.Z and H.Z critically reviewed the manuscript. All authors reviewed and approved the final version of the manuscript.

## Methods

### Data collection and processing

Antibody, nanobody and scFv entries targeting coronaviruses (CoVs) were obtained from CoV-AbDab (Raybould et al. 2021). Entries targeting other pathogens were collected through keyword-based searches in PubMed, bioRxiv, GenBank (Benson et al. 2011), and Google Patents, using term such as “RSV/hMPV/Influenza antibody”, “RSV/hMPV/Influenza antibody profiling”, “RSV/hMPV/Influenza B cell response”. A comprehensive set of references and patents was integrated into the database. For antibodies with resolved structures, we retrieved the structures from the Protein Data Bank (https://www.rcsb.org/) (Berman et al. 2000) by searching relevant keywords in the structure title. Therapeutic antibodies were collected from Thera-SAbDab (Dunbar et al. 2014) and PubMed. We additionally identified antibodies in clinical development by searching PubMed with the keyword “clinical trial” to capture candidates evaluated in clinical studies and to record their clinical stages.

All antibody amino acid and nucleotide sequences were obtained either directly from the original publications and patents or via GenBank using accession IDs cited in the source literature. Full-length variable region sequences were processed using ANARCI (Dunbar and Deane 2016) to annotate CDRs and assign corresponding V/J germline genes, which employs Hidden Markov Models to align input sequences to pre-numbered germline references. CDRs were annotated based on three standardized numbering schemes: the international IMGT (Lefranc et al. 2003), Kabat (Kabat and Wu 1971) and Chothia (Chothia and Lesk 1987).

### Viral genome sequence collection

Viral genome sequence data were collected from Nextstrain (https://nextstrain.org/) (Hadfield et al. 2018) and the NCBI database (NCBI Resource Coordinators 2017). Preliminary sample metadata were first downloaded from Nextstrain, and the strain names were extracted to facilitate sequence retrieval from NCBI. To minimize potential data loss, the NCBI Datasets command-line interface (CLI) was additionally used for direct sequence acquisition. Genome sequences were retrieved using taxon-specific keyword queries. For example, MERS-CoV genomes were obtained with the command: *datasets download virus genome taxon "Middle East respiratory syndrome coronavirus"*. Subsequently, the associated metadata were extracted using the command: *dataformat tsv virus-genome*. These procedures were applied for each target virus to generate a standardized dataset of genome sequences and corresponding metadata for downstream analyses.

### Antigen-antibody complexes interface residue analysis

For each antigen-antibody complex, the antigen chains and the corresponding antibody heavy and light chains were obtained from SAbDab (Dunbar et al. 2014). To characterize antigen-antibody interfaces, an in-house Python script was used to identify interface residues. Interface residues were defined as residues on the antigen and antibody partner chains with any interatomic distance of less than 4.5 Å. Each interface residue was annotated at the site level with entropy and recorded amino acid substitutions relative to a reference sequence.

### Normalized Shannon entropy analysis

After data collection, BLAST (v2.16.0) (Altschul et al. 1990) was used to compare each viral sequence against its corresponding reference genome in order to filter out low-quality sequences. Sequences were filtered using two criteria: (i) only alignments covering ≥ 80% of the reference genome were retained, and (ii) sequences with BLAST identity ≥ 80% were kept. The reference genomes used for each virus were: MERS-CoV (MF598664), SARS-CoV (NC_004718), SARS-CoV-2 (MN908947), RSV type A (PP109421), RSV type B (OP975389), hMPV type A (NC_039199) and hMPV type B (AY525843). The reference HA genes used for influenza virus were: h1n1 (AFM72832), h3n2 (AHG96407), h5n1 (AAD51927), h7n9(AHK10800), Victoria strain (ANC28539) and Yamagata strain (AET22022). Sequences passing these filters were subjected to multiple sequence alignment using MAFFT (v7.525) (Katoh et al. 2002). To quantify sequence variability at each position, we calculated the normalized Shannon entropy:*H_i_* = - ∑_*aϵA*_ *P_i_* (a) ln *P_i_* (a) where *H_i_* is the entropy at alignment position *i*, *A* denotes the set of amino acids observed at that position, and *p_i_ (a)* is the relative frequency of amino acid *a,* defined as *p_i_ (a) = (number of sequences containing residue a at position i) / (total number of sequences at position i)*. This formulation accounts for all possible residues observed at a given site.

### Mutation per-site analysis

Mutation frequencies were computed based on multiple sequence alignments. For each alignment position *i*, the reference residue was recorded, and the occurrences of all residues (including the reference) at that position were counted across all sequences. Gaps were excluded from the counts. For each observed residue *a*, the mutation frequency was defined as 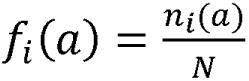 here *n_i_(a)* is the number of sequences containing residue *a* at position *i*, and *N* is the total number of sequences in the alignment.

### Phylogenetic clustering of antibody sequences

For a given set of antibody heavy chain nucleotide sequences, we first applied ANARCI (IMGT scheme) (Dunbar and Deane 2016) to extract the CDR3 region. To enable phylogenetic clustering, the antibody database was filtered to retain only entries containing nucleotide information and having the same CDR3 length as the input sequence. Sequences passing these filters were processed with Change-O (v1.3.3) (Gupta et al. 2015) and use hamming distance to define clonal groups. For each clone, all assigned sequences were aligned using MAFFT (v7.525) to generate multiple sequence alignments. These alignments were subsequently used as input for IQ-TREE (v2.2.2.6) (Nguyen et al. 2015). IQ-TREE was instructed to perform an extended model selection using ModelFinder Plus (MFP), invoked with the option -m MFP, which automatically identifies the most appropriate substitution model for the data and constructs the tree under that model. The resulting phylogenetic trees were further processed and visualized using ETE3 (Huerta-Cepas et al. 2016), a Python package for tree manipulation and visualization. In addition to tree construction, mutations were extracted from each internal node to characterize evolutionary changes along the branches. In addition, functionally characterized and uncharacterized sequences in the database were combined and pre-clustered using the same phylogenetic clustering strategy described above.

### Database implementation

The database was constructed using a modular, three-tier web architecture to enable efficient curation, querying, and visualization of antibody sequence variants and their functional implications. The backend is implemented with Spring Boot 3.4.4, providing a robust foundation for server-side logic, RESTful API design, and request handling through JavaServlets. Dynamic content rendering is achieved using the Thymeleaf template engine, which integrates seamlessly with HTML-based views to deliver data-driven pages. All curated antibody data including variable domain sequences (VH and VL), germline gene assignments, complementarity-determining regions (CDRs), somatic hypermutations, and experimentally validated functional variants are stored in a structured MySQL relational database. The database schema is normalized to minimize redundancy and ensure data consistency, with indexed fields on key identifiers such as antibody names, antigen targets, and mutation positions to optimize search performance. MyBatis is employed as the persistence layer to map Java objects to database records, supporting flexible and high-performance querying for complex use cases such as CDR-based filtering, mutation co-occurrence analysis, and lineage tracking. The frontend interface is built with HTML5, CSS3, and JavaScript, and enhanced with jQuery 3.6.0 for DOM manipulation, Semantic UI 2.5.0 for responsive and accessible user interface components, and DataTables 1.11.5 for interactive tabular display with sorting, pagination, and keyword search. To support structural interpretation of antibody–antigen interactions, the database integrates PDBe Molstar 3.3.0, a WebGL-powered molecular viewer, enabling users to visualize the 3D structure of Fab or Fv fragments and explore the spatial context of mutations within antigen-binding sites directly within the browser.

