## Supplemental Figure1-6 and Table 1 for "MAAD: Multidimensional Antiviral Antibody Database": SupplementaryMaterials.docx

**List of Supplementary Materials**

Figure S1. Overview of entries in MAAD.

Figure S2. CDR3 length distributions of entries in MAAD.

Figure S3. Circos plots of germline gene pairings of antibodies.

Figure S4. Detail page of MAAD showing

Figure S5. Viral entropy and mutation analysis.

Figure S6. Workflow of sequence-based clustering and tree construction.

Table S1. Summary of fields in the MAAD database

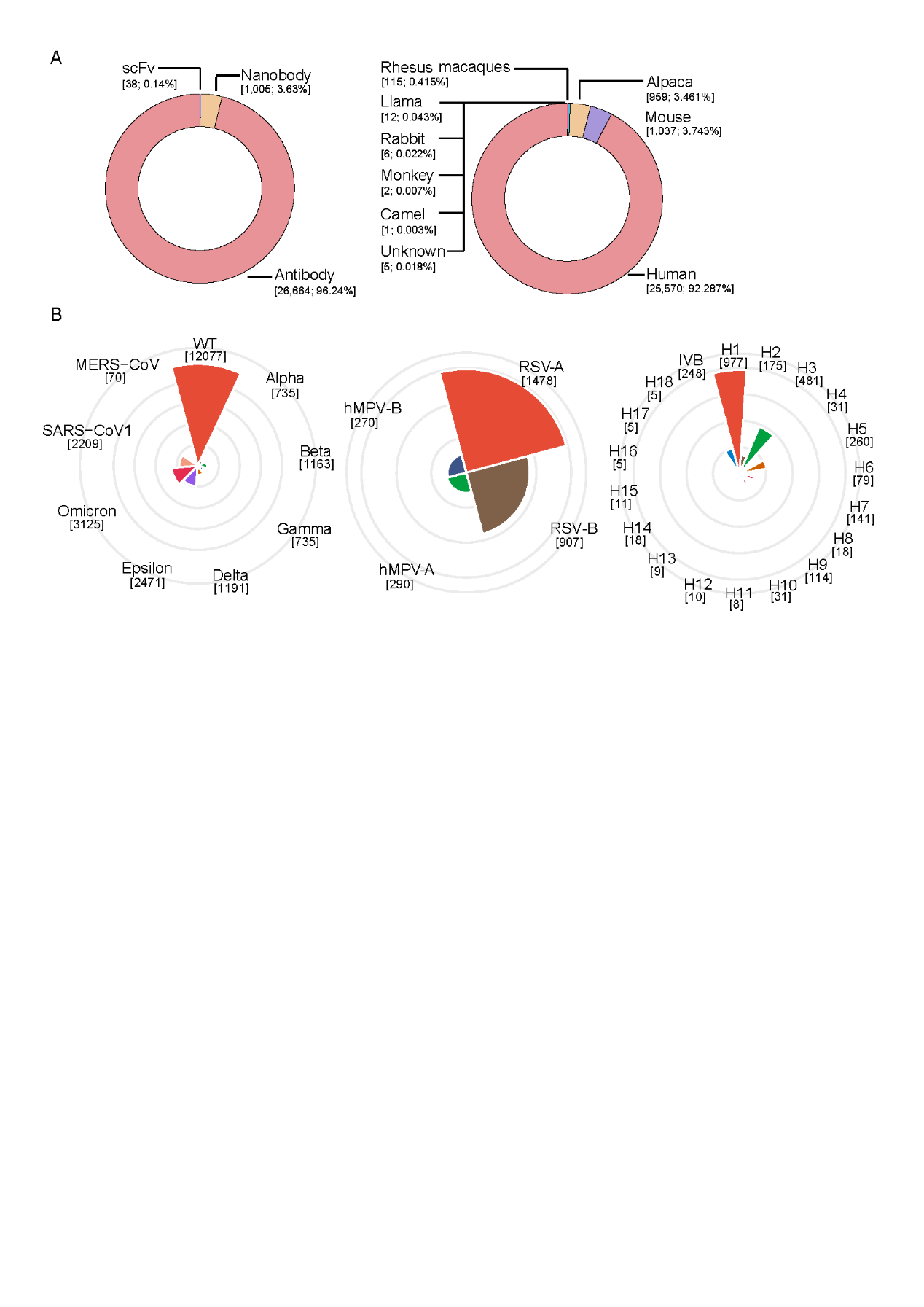

**Figure S1. Overview of entries in MAAD.**

(A) Donut plots showing the distribution of antibodies, nanobodies, and scFv (left); and the developmental species origin of each entry in MAAD (right). (B) Polar bar plot summarizing the number of antibodies, nanobodies and scFvs annotated to bind to or neutralize each viral strain/subtype. *Coronaviridae*: SARS-CoV-2 (WT, Alpha, Beta, Gamma, Delta, Epsilon, Omicron), SARS-CoV-1 and MERS-CoV (right). *Pneumoviridae*: RSV-A, RSV-B, hMPV-A, and hMPV-B (middle). *Orthomyxoviridae*: Influenza A H1–H18 and influenza B (right). Bar lengths encode counts; labels indicate the exact number of unique antibodies, nanobodies and scFvs per category. Colors distinguish categories only.

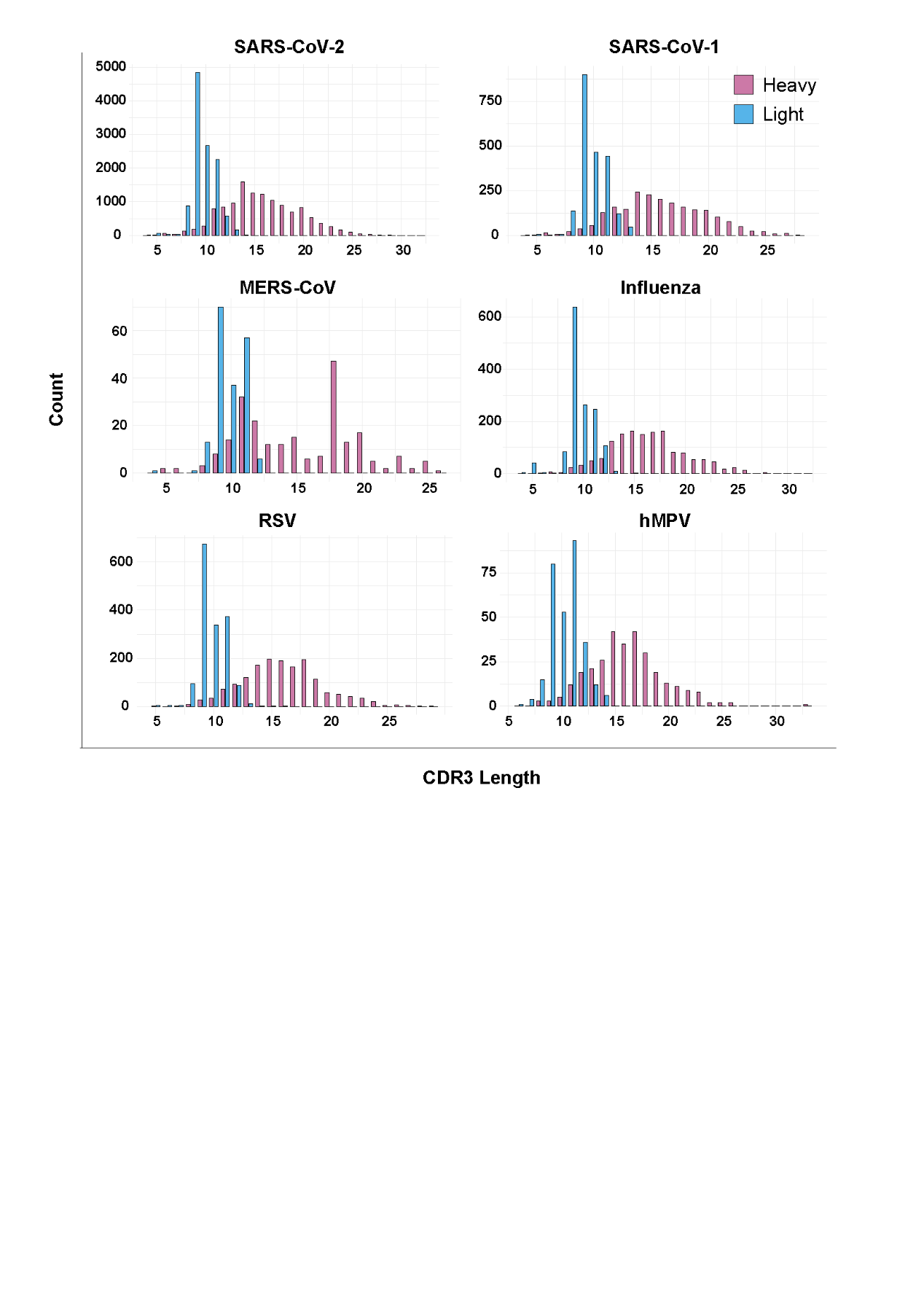

**Figure S2. CDR3 length distributions of entries in MAAD.**

Bar plots show the distributions of heavy-chain (pink) and light-chain (blue) CDR3 lengths numbered by IMGT scheme of antibodies, nanobodies and scFvs.

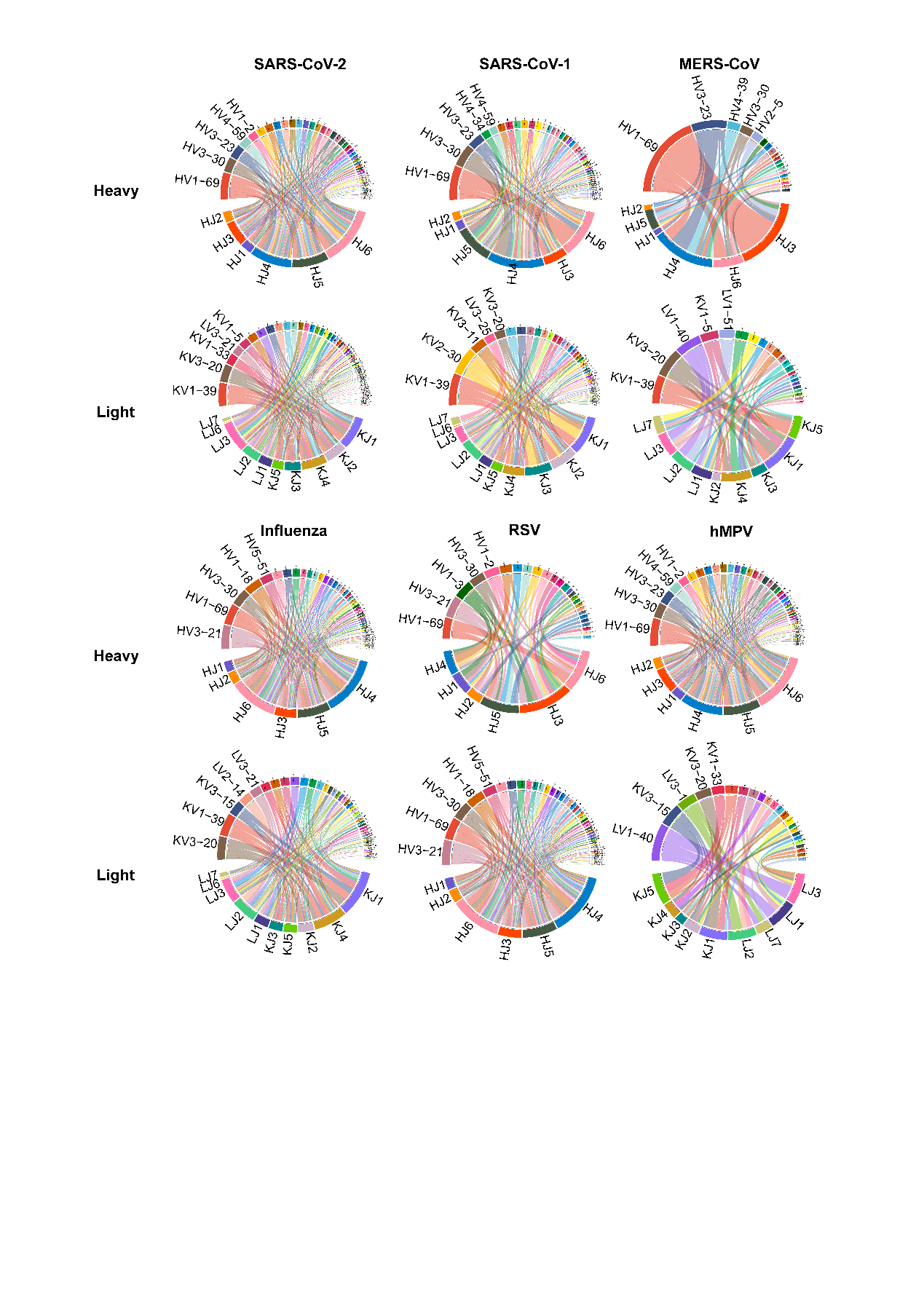

**Figure S3. Circos plots of germline gene pairings of antibodies.**

Circos plots showing IGHV-IGHJ (heavy chain) and IGLV-IGLJ (light chain) gene pairings observed in antibodies targeting SARS-CoV-2, SARS-CoV-1, MERS-CoV, influenza, RSV, and hMPV. Each arc represents a germline gene segment, and connecting ribbons indicate observed pairings between V and J genes. The width of the ribbons reflects the frequency of each pairing within the dataset.

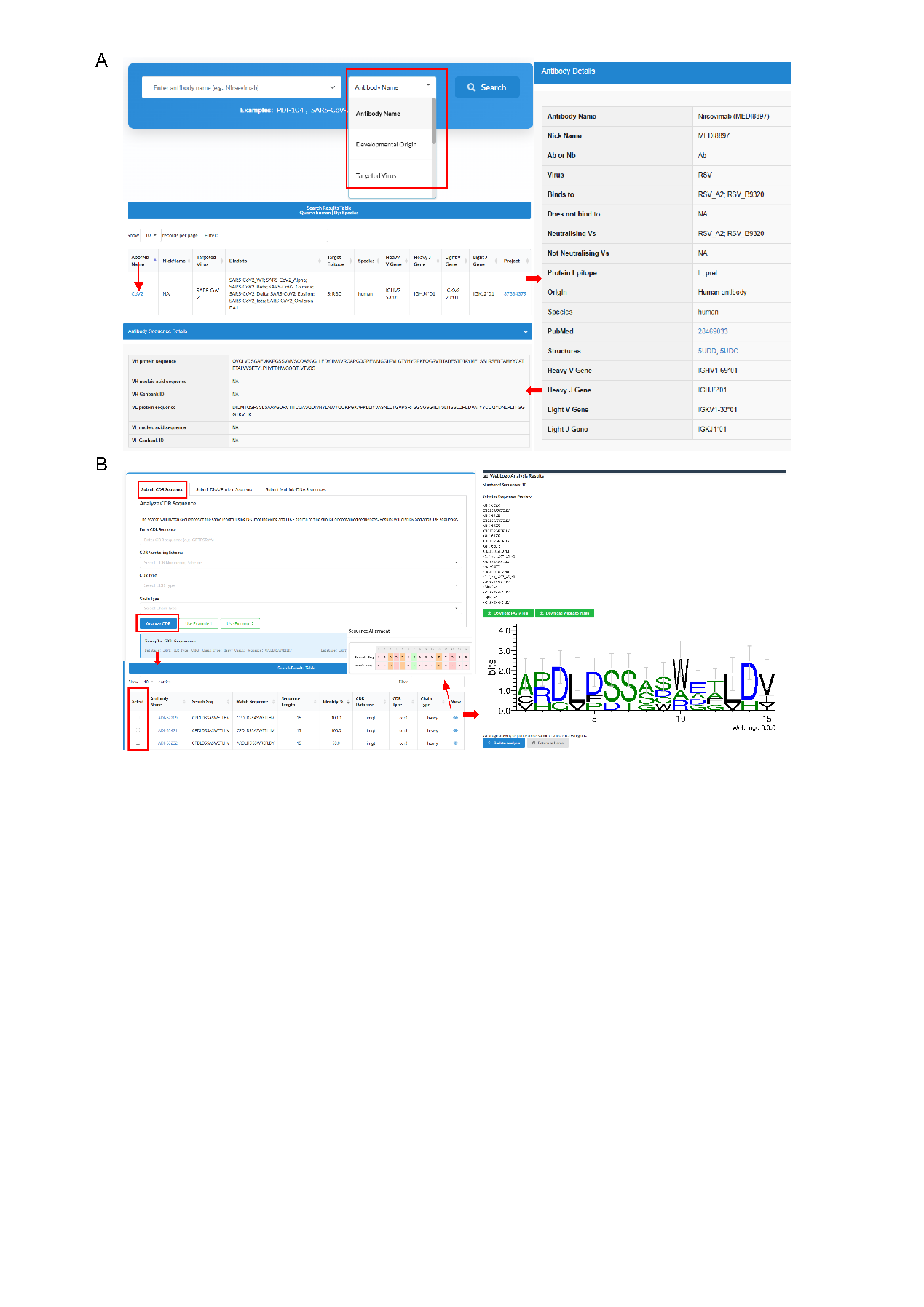

**Figure S4. Detail page of MAAD showing**

(A) Search interface for querying entries and viewing detailed information, including binding/neutralization profiles, sequences, germline genes, and CDRs. (B) Interactive interface and CDR-focused analysis results with selected CDR logo plots.

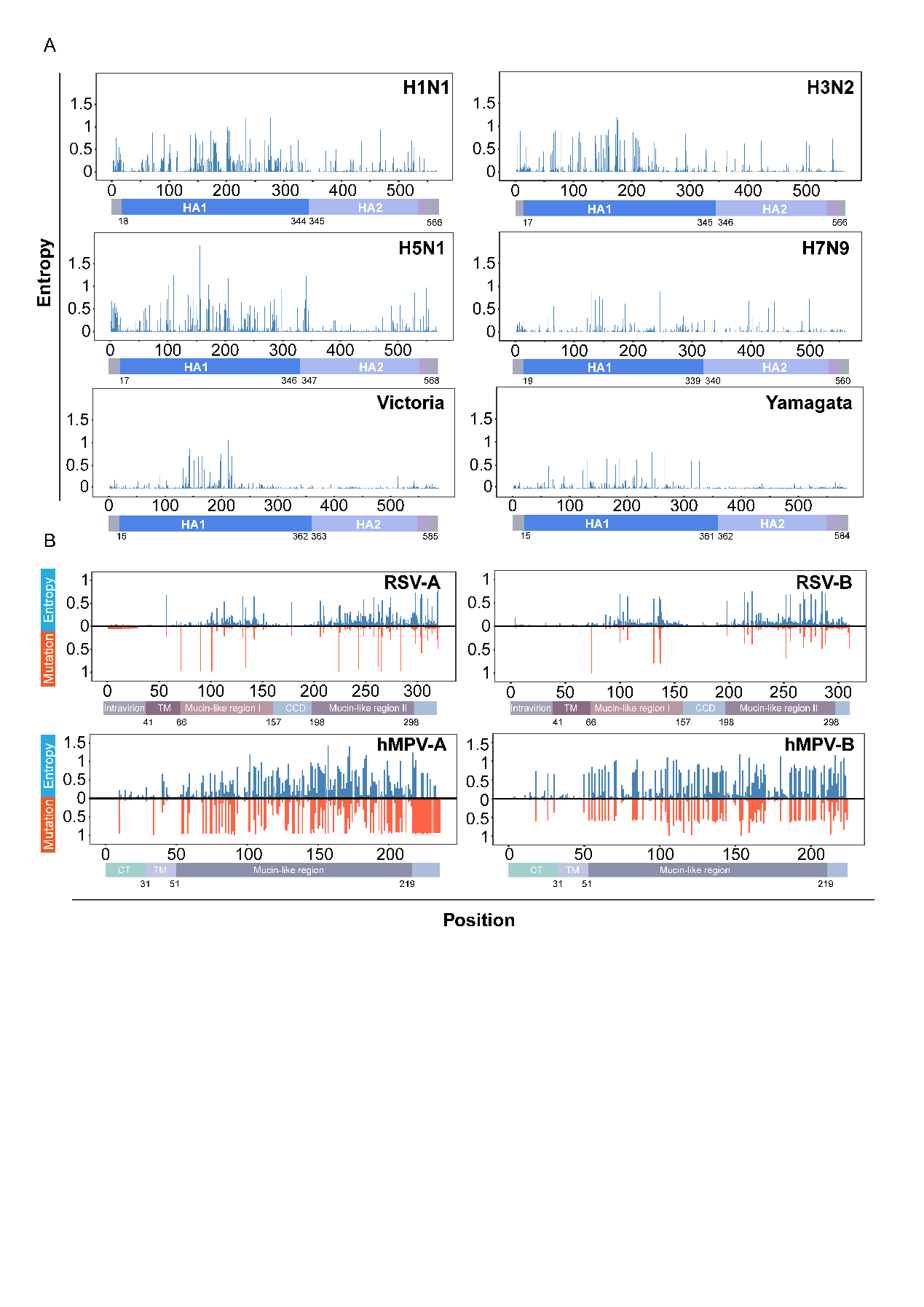

**Figure S5. Viral entropy and mutation analysis.**

(A) Site-specific Shannon entropy calculated from the aligned HA sequences from H1N1, H3N2, H5N1, and H7N9 and influenza B virus (Victoria and Yamagata lineages) to quantify variability. (B) Bar plots display per-residue Shannon entropy (blue) and mutation frequency (orange) of RSV and hMPV G protein, calculated based on aligned viral sequences relative to a reference strain

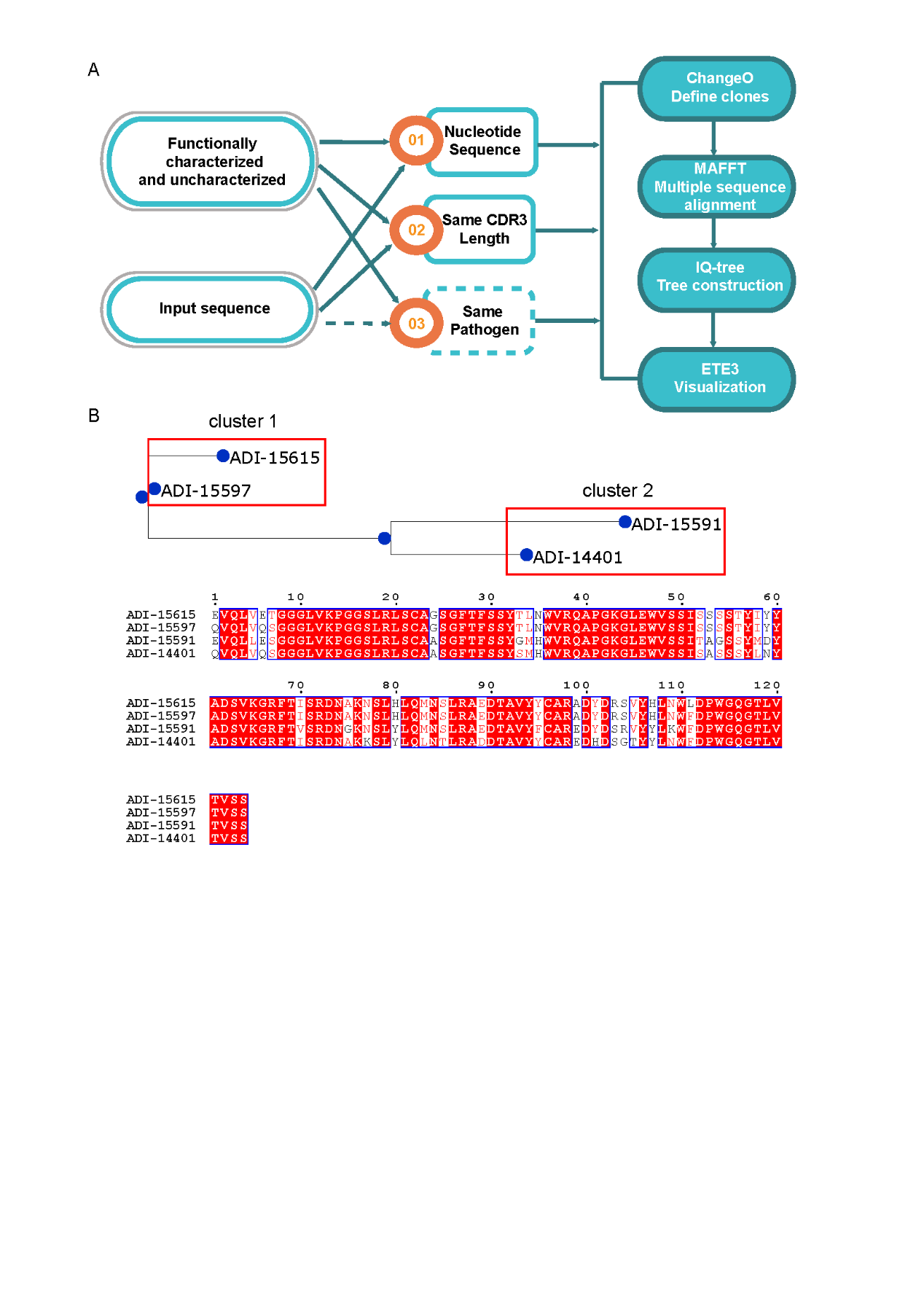

**Figure S6. Workflow of sequence-based clustering and tree construction.**

(A) MAAD supports two modes of clustering analysis: (1) exploration of precomputed phylogenetic trees that incorporate both functionally characterized and uncharacterized entries across the database and (2) user-driven phylogenetic tree reconstruction, in which user-uploaded sequences are integrated with MAAD sequences to infer a combined phylogenetic tree. In user-driven modes, nucleotide sequences are first grouped by CDR3 length and pathogen source (optional). In both modes, clonal assignment is performed with Change-O, multiple sequence alignment is conducted using MAFFT, and phylogenetic trees are constructed with IQ-TREE and visualized using ETE3. (B) An example of phylogenetic tree and corresponding multiple sequence alignment.

| Table S1. Summary of fields in the MAAD database | |
| --- | --- |
| Seq | Unique sequence identifier for Ab, Nb and scFv |
| Name | Published name of Ab, Nb and scFv |
| Nickname | Commonly used nickname of Ab, Nb and scFv |
| AborNb | Specifies whether the entry is an Ab, Nb or scFv |
| Virus | The virus from which B cells producing the Ab, Nb or scFv were derived |
| Bindsto | Antigens experimentally confirmed to bind to Ab, Nb and scFv |
| Doesnot Bindto | Antigens experimentally confirmed not to bind to Ab, Nb and scFv |
| NeutralisingVs | Antigens experimentally neutralized by Ab, Nb and scFv |
| Not NeutralisingVs | Antigens experimentally tested but not neutralized by Ab, Nb and scFv |
| Protein Epitope | Protein domain targeted by Ab, Nb and scFv |
| Origin | Developmental biological or synthetic origin of Ab, Nb and scFv (e.g. human, murine, engineered) |
| Species | Species of origin of Ab, Nb and scFv |
| VHorVHH | Heavy variable domain amino acid sequence |
| VH nuc | Heavy variable domain nucleotide sequence |
| VHGenbankID | Accession number of GenBank for the heavy variable domain nucleotide sequence |
| VL | Light variable domain amino acid sequence |
| VL nuc | Light variable domain nucleotide sequence |
| VLGenbankID | Accession number of GenBank for the light variable domain nucleotide sequence |
| Structures | Links to available antigen–antibody/nanobody complex structures |
| PMID | PubMed identifier linking to the primary reference |
| Reference | References to the primary literature on Ab, Nb and scFv |
| Last updated | Timestamp indicating when the Ab, Nb and scFv was added |
| Pub date | Publication date of Ab, Nb and scFv |
| Heavy/Light identity species | Species of the most identical germline heavy or light chain |
| Heavy/Light V Gene | The identifier of the most sequence identical germline over the heavy or light chain v-region |
| Heavy/Light J Gene | The identifier of the most sequence identical germline over the heavy or light chain j-region |
| CDRH/L1, CDRH/L2, CDRH/L3 (imgt, kabat, chothia) | IMGT, kabat and chothia numbering for heavy and light chain CDRs |
| CDRH/L1_len, CDRH/L2_len, CDRH/L3_len (imgt, kabat, chothia) | CDR lengths of heavy and light chain (IMGT, kabat and chothia scheme) |
